# Continued circulation, recombination and evolution of the ancient subcontinent lineage despite predominance of the recent arctic-like lineage rabies viruses (RABV) in India

**DOI:** 10.1101/398180

**Authors:** Jagadeeswaran Deventhiran, Bhuvaneswari Srinivasan, Manoharan Seeralan, Vijayarani Kanagaraj, Sangeetha Raju, Kumanan Kathaperumal, Ruth Nissly, Sunitha Manjari Kasibhatla, Mohan M Kale, Urmila Kulkarni-Kale, Suresh V Kuchipudi

## Abstract

**Background:** Rabies is an emerging and re-emerging lethal encephalitis causing 26,400 to 61,000 human deaths annually. Approximately 20,000 people die of rabies every year in India that accounts to 36% of the world’s rabies deaths. Rabies is endemic among domestic dogs in India and there are conflicting reports on the currently circulating RABV lineages in domestic dogs in India. Further, movement of humans and animals between Sri Lanka and southern coastal states of India was proposed to be a source of the emergence of variant RABV in India. For effective prevention and control of rabies in India it is essential to establish the genetic diversity and evolutionary dynamics of RBAV currently circulating in India.

**Methods:** We carried out molecular evolution and recombination analyses of nucleoprotein (N) and glycoprotein (G) genes of 26 RABV isolates from southern Indian states of Tamil Nadu and Goa.

**Results:** We found continued co-circulation of ancient subcontinent lineage despite predominance of the recent arctic-like lineage RABVs in southern India. The mean rate of nucleotide substitution in G and N genes was 1.32 × 10^−3^ and 1.91 × 10^−4^ substitutions/site/yr respectively. The study also found recombination in both N and G genes and a higher mean rate of evolutionary changes in G gene among Indian dog RABV isolates than those of lyssaviruses. The Indian subcontinent lineage RABV isolates investigated in this study clustered closely with other subcontinent lineage viruses from Sri Lanka highlighting the continued incursion and/or circulation of the variant subcontinent lineages of RABVs between India and Sri Lanka.

**Conclusion:** We report that there is enzootic viral establishment of two distinct RABV lineages in domestic dogs in India that are evolving at a greater rate.

**Author summary:** Rabies is a fatal viral disease that has no treatment and can only be prevented by post-exposure vaccination. In many parts of Asia and Africa, rabies continues to be a major public health threat almost always caused by dog bites. In this study, we investigated the genetic diversity and rate of evolution among rabies viruses isolated from dogs in India. We found that two distinct lineages of Rabies viruses (RABVs) namely the ancient subcontinent lineage and a more recent arctic-like lineage co-circulate among dogs in India. Notably, our study found that the dog rabies viruses in India are undergoing recombination and evolving at a higher rate than other lyssaviruses. Phylogenetic analysis revealed continued incursion and/or circulation of the variant subcontinent lineages of RABVs in India that might have been originated from Sri Lanka. Our study indicates that two distinct lineages of RABVs are maintained and currently circulate among dog population in India

## Introduction

Rabies is one of the longest known human diseases and was first documented at least 4,000 years ago [1-3]. Rabies is among the key neglected tropical diseases (NTDs) that predominantly affects poor and vulnerable populations. According to World Health Organization (WHO) estimates in 2010, the annual number of human deaths due to rabies globally ranges between 26,400 to 61,000 with the vast majority of deaths (84%) occurring in rural areas [4].

Rabies is caused by Rabies virus (RABV), which belongs to the genus *Lyssavirus*, belonging to the family *Rhabdoviridae* and order Mononegavirales [5]. RABV virions are enveloped, rod- or bullet-shaped, with single stranded negative sense RNA genomes measuring approximately 12 kb. Among the 12-identified species within the genus *Lyssavirus*, RABV has the broadest geographic distribution and the widest spectrum of vectors or reservoir hosts within the orders Carnivora and Chiroptera [6-8]. RABV is the main etiological agent of rabies, an acute and almost invariably fatal form of encephalomyelitis, which can affect almost all terrestrial mammals, including humans. Contamination with infected saliva by bite, scratch or mucous membrane exposure constitutes an important source of infection. Despite the availability of an effective post-exposure prophylaxis, it is estimated that approximately 55,000 people die every year due to rabies, with more than 95% of human deaths occurring in Asia and Africa [9]. According to 2011 WHO estimates, 1.74 million disability-adjusted life years are lost each year due to rabies and estimated annual cost burden is US$ 583.5 million. Though rabies can be prevented by vaccination, there is no effective treatment after the onset of the clinical disease [6].

The domestic dog remains the main reservoir and vector of rabies in developing countries and is responsible for almost all human deaths. Dogs have been identified as the main vector involved in interspecies RABV transmission. Various wild carnivores are involved in the maintenance of RABV and transmission of sylvatic rabies in limited geographic regions, with a small contribution in the burden of human rabies. Other terrestrial mammal species including livestock species are susceptible to rabies but act as epidemiological dead-end hosts as they do not transmit the disease further [10]. Multiple lineages of dog RABVs have been described that include African, Asian, Arctic-like, Cosmopolitan and Indian subcontinent [3]

Rabies continues to be major public health threat in India. A national rabies survey in India in 2006 sponsored by the WHO reported that 20,000 persons died of rabies each year [11]. Molecular epidemiology of RABV isolates in India based on nucleoprotein (N) and phosphoprotein (P) genes found that all the Indian RABV isolates clustered within a single clade corresponding to Arctic/Arctic-like viruses [12]. Using the ecto-domain coding region of the glycoprotein gene sequence, a study later found co-existence of both Arctic like 1 lineage and Indian subcontinent lineage RABV in India [13]. However, a more recent study reported that based on G gene sequence, all the Indian isolates obtained between 2001-2014 from six different species of animals were genetically related to Arctic-like 1a lineage viruses [14]. Indian RABV isolates clustered within the Arctic/Arctic-like clade are well separated in evolutionary terms from the Cosmopolitan lineage as well as other lineages that circulate in various parts of Southeast Asia [15]. Most of the RABV molecular epidemiological studies carried out in India were done with isolates collected over a long period of time and from multiple geographical locations in India. To design better prevention and control strategies, it is important to investigate the genetic diversity and evolutionary dynamics of RABV lineages.

Molecular epidemiologic approaches are very helpful to track the spread of RABV variants and identify their incursion into new geographic regions [16]. Further, molecular epidemiological analyses of RABVs in Rabies endemic countries such as India would allow accurate analysis of the spread and evolution of RABVS to design better control methods [15]. A previous study found a RABV isolate from the city of Chennai (previously known as Madras) in the southeastern coast in India clustered more closely to a distinct variant RABV found in Sri Lanka rather than with the other Indian isolates [15]. Chennai, in the state of Tamil Nadu, is the southernmost city in India. The movement of humans and their animals between Sri Lanka and India was proposed to be a source of the emergence of the variant RABV in India [15]. Movement of people continues to occur between the southeastern coastal area of India and Sri Lanka. In order to better understand the genetic divergence of RABV lineages circulating in India, we carried out molecular evolutionary analyses of RABV isolates collected in 2014 from southern Indian states of Tamil Nadu and Goa.

## Methods

### Samples and laboratory tests to confirm rabies

Twenty-six brain samples from animals, which died of rabies symptoms, were used in this study. The samples included twenty-three isolates from domestic dogs, two samples from goats, and one sample from cattle. The samples were collected in 2014 from two different states on the east and west coast of southern India (Tamil Nadu and Goa) (Table 1). The samples were tested at the rabies unit of Madras Veterinary College for rabies confirmation by fluorescent antibody test (FAT) [17]. In addition to FAT, N-gene specific RT-PCR [18] of 20% homogenate of the respective brain samples in phosphate buffered saline was performed to confirm the presence of rabies viral genomes. Sample collection from animals was performed in full compliance with the Committee for the Purpose of Control and Supervision of Experiments on Animals (CPCSEA) regulations and the study was approved by the Institutional Animal Ethics Committee (IAEC) of Tamil Nadu Veterinary and Animal Sciences University (TANUVAS), Chennai, India.

**Table 1:**
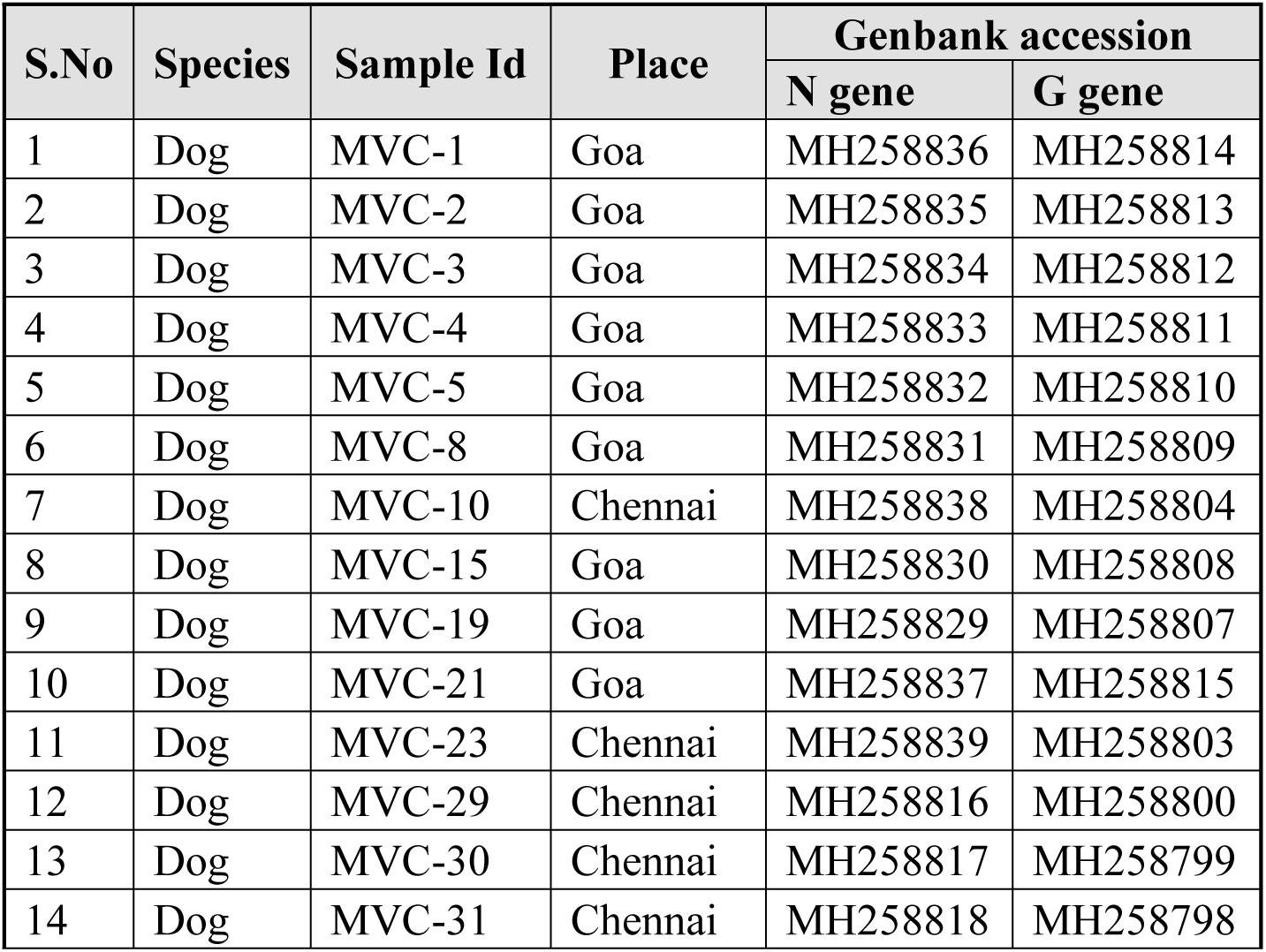
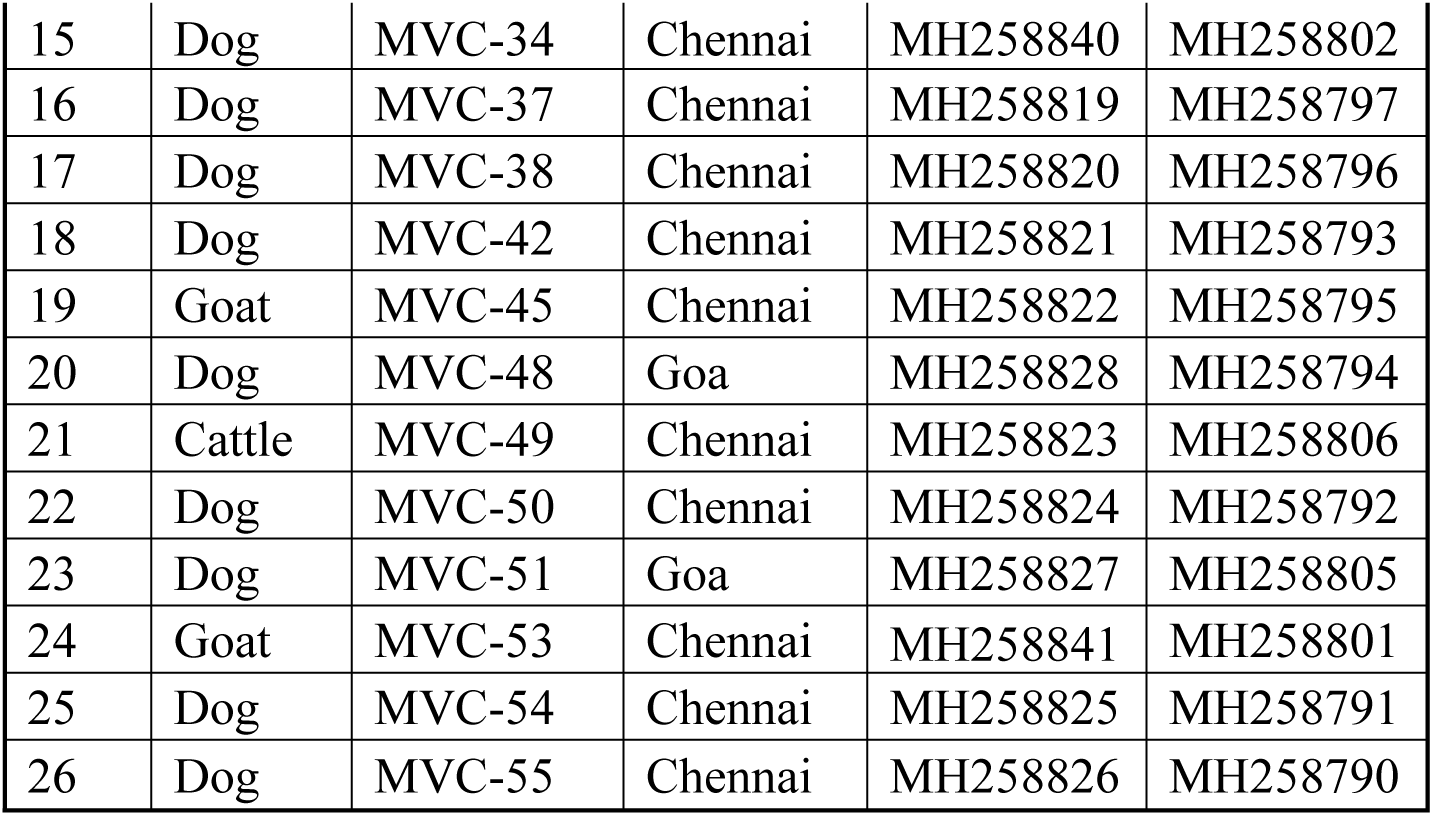
List of rabies positive samples and their origin

Samples and laboratory tests to confirm rabies:
| S.No | Species | Sample Id | Place | Genbank accession |  |
| --- | --- | --- | --- | --- | --- |
|  |  |  |  | N gene | G gene |
| 1 | Dog | MVC-1 | Goa | MH258836 | MH258814 |
| 2 | Dog | MVC-2 | Goa | MH258835 | MH258813 |
| 3 | Dog | MVC-3 | Goa | MH258834 | MH258812 |
| 4 | Dog | MVC-4 | Goa | MH258833 | MH258811 |
| 5 | Dog | MVC-5 | Goa | MH258832 | MH258810 |
| 6 | Dog | MVC-8 | Goa | MH258831 | MH258809 |
| 7 | Dog | MVC-10 | Chennai | MH258838 | MH258804 |
| 8 | Dog | MVC-15 | Goa | MH258830 | MH258808 |
| 9 | Dog | MVC-19 | Goa | MH258829 | MH258807 |
| 10 | Dog | MVC-21 | Goa | MH258837 | MH258815 |
| 11 | Dog | MVC-23 | Chennai | MH258839 | MH258803 |
| 12 | Dog | MVC-29 | Chennai | MH258816 | MH258800 |
| 13 | Dog | MVC-30 | Chennai | MH258817 | MH258799 |
| 14 | Dog | MVC-31 | Chennai | MH258818 | MH258798 |

**Table 1:** List of rabies positive samples and their origin
|  |  |  |  |  |  |
| --- | --- | --- | --- | --- | --- |
| 15 | Dog | MVC-34 | Chennai | MH258840 | MH258802 |
| 16 | Dog | MVC-37 | Chennai | MH258819 | MH258797 |
| 17 | Dog | MVC-38 | Chennai | MH258820 | MH258796 |
| 18 | Dog | MVC-42 | Chennai | MH258821 | MH258793 |
| 19 | Goat | MVC-45 | Chennai | MH258822 | MH258795 |
| 20 | Dog | MVC-48 | Goa | MH258828 | MH258794 |
| 21 | Cattle | MVC-49 | Chennai | MH258823 | MH258806 |
| 22 | Dog | MVC-50 | Chennai | MH258824 | MH258792 |
| 23 | Dog | MVC-51 | Goa | MH258827 | MH258805 |
| 24 | Goat | MVC-53 | Chennai | MH258841 | MH258801 |
| 25 | Dog | MVC-54 | Chennai | MH258825 | MH258791 |
| 26 | Dog | MVC-55 | Chennai | MH258826 | MH258790 |

### RNA extraction and RT-PCR

Total RNA was extracted from the brain tissue homogenates using TRIzol reagent (Invitrogen), following the manufacturer’s instructions. N and G gene specific primers were designed based on the available rabies sequences of the Indian isolates (Table 2). G gene was amplified by RT-PCRs as two overlapping fragments. Two-step RT-PCR was performed using 1μg of total RNA. Complementary DNA (cDNA) synthesis was done using Superscript Reverse Transcriptase III enzyme (Invitrogen) per manufacturer’s instruction. PCR amplification was done using Platinum Pfx polymerase (Invitrogen) and thermal cycler profile was as follows: polymerase activation at 94 °C for 5 min; 35 cycles of denaturation at 94 °C for 15 s, annealing at 55 °C for 30 s, and extension at 68 °C for 1 min; extension at 72 °C for 10 min.

**Table 2:**
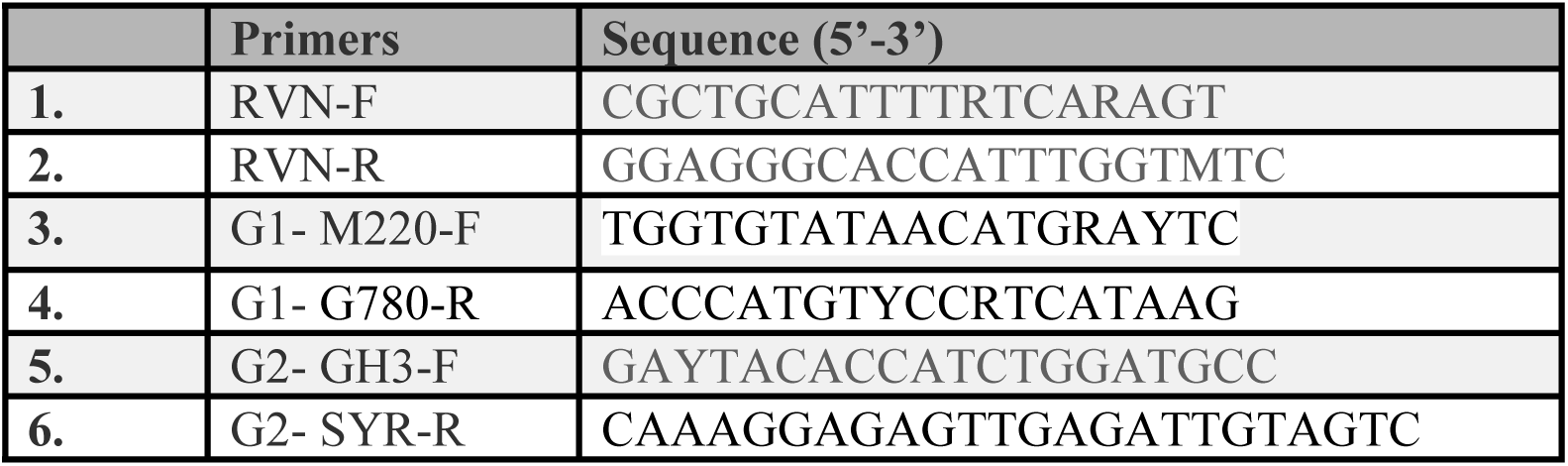
Gene specific primers used for gene amplification and sequencing.

### Sequencing

Amplified gene segments were purified by QIAquick gel purification kit (Qiagen) per the manufacturer’s instructions. DNA sequencing of the N and G genes were performed using the Illumina HiSeq2000 platform (Illumina). For 26 isolates of RABV, complete nucleotide sequencing of nucleoprotein (N) gene (1353 nucleotides) and partial sequencing of glycoprotein (G) gene (711 nucleotides) was performed.

### Phylogenetic & evolutionary analysis

Multiple sequence alignment was carried out using MUSCLE [19, 20]. Recombination detection was carried out using RDP4 [21]. Phylogenetic reconstruction was carried out using both alignment-based and alignment-free methods. An alignment-based phylogenetic tree was derived using the maximum likelihood (ML) method in the PhyML package [22] with 1000 bootstrap replicates. An alignment-free method based on Return Time Distribution (RTD) developed in house was used with k-mer value of 6 to derive trees [23].

Molecular clock analysis was carried out using BEAST v1.8.4 [24]. Earlier model selection was carried out using jModelTest [25] and bat rabies virus was used as an outgroup. Relaxed clock with lognormal distribution was used for molecular clock analysis with GTR+I+gamma as substitution model and constant coalescence as demographic model. Markov chain Monte Carlo (MCMC) algorithm was run for 6 million steps and sampled every 1000 steps. Tracer v1.7 [26] was used for assessing convergence, and iTOL server was used for visualization of the phylogenetic trees [27].

## Results

### Phylogenetic and evolutionary analysis based on N gene

Sequencing of complete nucleoprotein gene of 26 Indian isolates (length 1353 nucleotides) was carried out. Sequence data of 15 isolates were used for further analysis based on removal of in-frame stop codons and detection of recombination events. Recombination analysis revealed that four isolates, namely MVC-21, MVC-34, MVC-50 and MVC-53 (GenBank accession numbers MH258837, MH258840, MH258824 and MH258841, respectively) are potential recombinants as predicted by three or more methods with p-value < 0.0005 and were hence excluded from molecular phylogenetic analysis.

The 15 nucleoprotein sequences of Indian isolates along with representative RABV sequences from across the globe (totaling 75 sequences) were used for molecular phylogenetic analysis (MPA) using both alignment-based and alignment-free methods. Multiple sequence alignment revealed that sequences shared ∼81% identity and ∼91% similarity. The maximum likelihood phylogenetic tree showed clustering based on known lineages, namely Arctic-like, Africa 2, Indian subcontinent, Cosmopolitan and Asian (Fig 1). The mean rate of nucleotide substitution of N gene nucleotides estimated using by a Bayesian method was 1.91 × 10^−4^ substitutions/site/yr (95% highest posterior density (HPD) 1.05 × 10^−4^ to 2.78 × 10^−4^). Based on the N gene, the RABV isolates sequenced in this study clustered into two lineages, Arctic-like (13 isolates) and Indian subcontinent (2 isolates). Tree topology generated using an alignment-free method (Fig 2) agreed with that of maximum likelihood and Bayesian methods. Arctic-like lineage RABV isolates reported in this study clustered closely with Arctic-like lineage RABVs from neighboring countries like Nepal and Afghanistan. The Indian subcontinent lineage RABV isolates reported in this study clustered with previously reported Indian subcontinent lineage RABV isolates from India and Sri Lanka.

**Fig 1:**
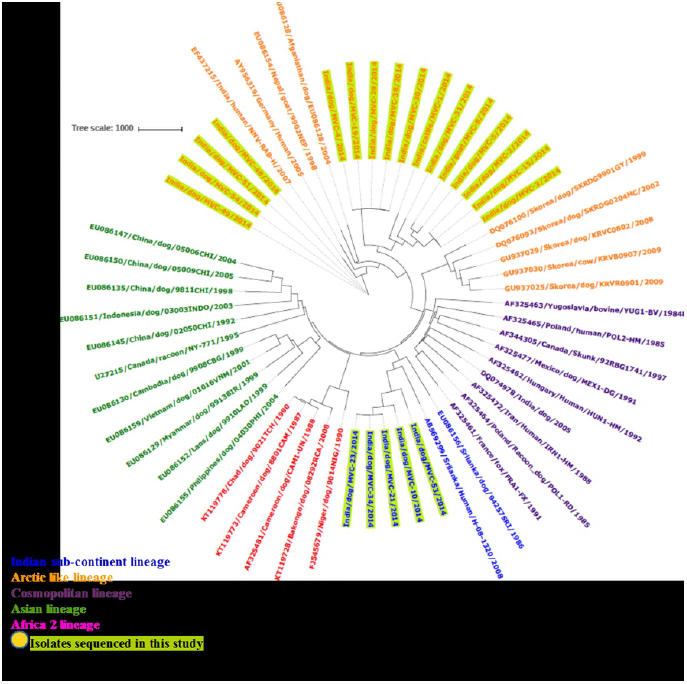
Maximum likelihood tree based on Nucleoprotein (N) gene. N gene sequences of 15 Rabies virus (RABV) Indian isolates along with representative RABV sequences from across the globe (totaling 75 sequences) were used to construct the Maximum likelihood tree using PhyML with 1000 bootstrap replicates.

**Fig 2:**
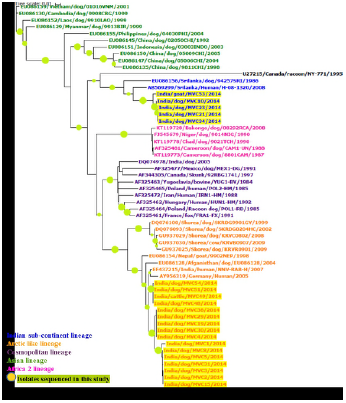
RTD-based alignment-free phylogenetic tree based on Nucleoprotein (N) gene. RTD-based alignment-free phylogenetic tree (derived using K-mer = 6) of nucleoprotein (N) sequences of 15 Rabies virus (RABV) Indian isolates along with representative RABV sequences from across the globe (totaling 75 sequences).

### Phylogenetic and evolutionary analysis based on G gene

Partial sequencing of RABV glycoprotein gene (711 nucleotides) was carried out for 26 Indian isolates. Sequence data of only 21 isolates were used for further analysis based on removal of sequences containing in-frame stop codons and detection of recombination events. Recombination event were detected in isolate MVC-50 (GenBank accession number MH258824) by more than three methods with p-value < 0.0005 and hence was removed from further phylogenetic analysis.

The 21 glycoprotein sequences of Indian isolates along with representative RABV sequences from across the globe (totaling 57 sequences) were used for molecular phylogenetic analysis using alignment-based and alignment-free methods. MSA showed ∼70% identity and ∼84% similarity. Lineages of the various RABV isolates were estimated using ML method implemented in PhyML package. The mean rate of nucleotide substitution estimated from the partial glycoprotein sequences by Bayesian analysis was 1.32 × 10^−3^ substitutions/site/yr (95% HPD 6.92 × 10^−4^ to 2.05 × 10^−3^).

Similar clustering patterns of RABV isolates were observed in the trees generated using alignment-based (ML, (Fig 3) and Bayesian) and alignment-free (RTD, Figure 4) methods. Of the Indian RABV isolates, 16 RABV clustered into Arctic-like lineage and 5 clustered into Indian subcontinent lineage. The phylogenetic trees of the N and G genes (Fig 1, 2, 3 and 4) displayed similar topologies, indicating the presence of equivalent clades in both trees. For example, both trees showed Indian subcontinent lineage RABV isolates clustering closely with other subcontinent Indian subcontinent lineage RABV isolates from Sri Lanka. Notably, maximum likelihood tree showed that the Arctic like RABV isolates from this study clustered closely with human RABV isolates from India and Germany.

**Fig 3:**
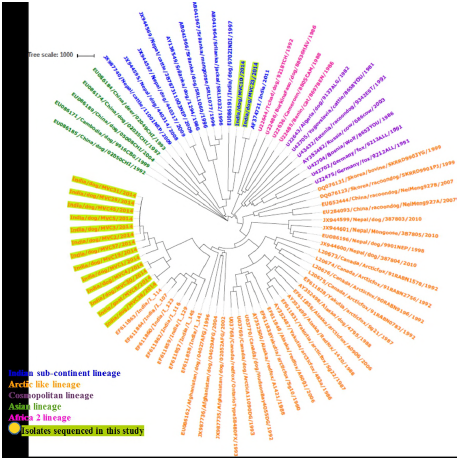
Maximum likelihood tree based on Glycoprotein (G) gene. G gene sequences of 21 Rabies virus (RABV) Indian isolates along with representative RABV sequences from across the globe (totaling 57 sequences) were used to construct the Maximum likelihood tree using PhyML with 1000 bootstrap replicates.

**Fig 4:**
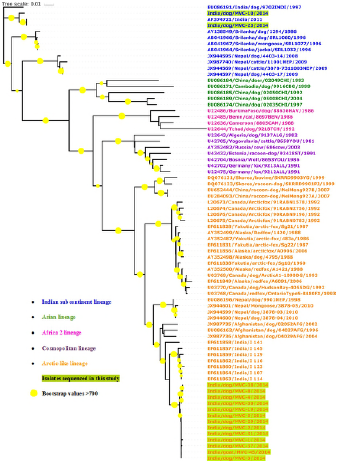
RTD-based alignment-free phylogenetic tree based on Glycoprotein (G) gene. RTD-based alignment-free phylogenetic tree (derived using K-mer = 6) of glycoprotein sequences of 21 Rabies virus (RABV) Indian isolates along with representative RABV sequences from across the globe (totaling 57 sequences).

The time-scaled evolutionary trees obtained using Bayesian analysis (data not shown) based on both N and G genes indicated that the isolates of Arctic-like lineage diversified more recently compared to isolates of Indian subcontinent lineage.

## Discussion

A key aspect of RNA viruses is their exceptionally variable rates of molecular evolution with up to 6 orders of magnitude in nucleotide substitution rates has been observed among viral species [28]. Like all RNA viruses, RABVs constantly evolve, and the rate of evolutionary change in different host species is determined by the nature of virus-host interactions [29]. We investigated the evolutionary rates of RABV isolates from India using partial sequences of the glycoprotein (G), the surface protein that allows RABV to enter the cells of nervous system, and the complete gene sequences of nucleoprotein (N), which forms the viral capsid and plays a role in transcription and replication.

The mean rate of nucleotide substitution estimated from the N gene sequences was 1.91 × 10^−4^ substitutions/site/yr (95% HPD 1.05 × 10^−4^ to 2.78 × 10^−4^). Interestingly, these estimates are in close agreement with the reported N-gene mean rate of substitution in RABVs from bats and terrestrial mammals [30] suggesting that the dog RABVs in India continue to evolve at the same rate as bat RABVs from around the world. The mean rate of nucleotide substitution estimated from the partial G gene sequences is 1.32 × 10^−3^ substitutions/site/yr (95% HPD 6.92 × 10^−4^ to 2.05 × 10^−3^) which is marginally higher than that reported in a previous study, which evaluated substitution rate in Indian RABV isolates based on the ecto domain of glycoprotein [13]. Earlier study using the complete G gene sequences reported the mean rate of substitution to be 3.9×10^−4^ subs per site per year; 95 % HPD values=1.2–6.5×10^−4^ subs per site per year [31]. The results of this study suggest a higher mean rate of evolutionary change in G gene in Indian dog RABV isolates as compared to evolutionary rates of lyssaviruses. The observed rate of evolution could be attributed to use of partial vs full G gene sequences for analysis and the number of isolates used for estimating the rate of evolution. Therefore, there is a need to re-estimate rate of substitution of G gene using a higher sample size covering both, Indian and global population of RABVs.

Recombination events have been reported in the polymerase gene of RABV using the complete genome data [32]. This study reports recombination in both N and G genes of Indian RABV isolates. Four Indian RABV isolates (MVC-21, MVC-34, MVC-50 and MVC-53) displayed sites for potential recombination events when analyzed using the N-gene sequences. Interestingly, MVC-50 was also predicted to be a potential recombinant using the G gene partial sequence. Homologous recombination is known to occur among rabies viruses which could play a role in the diversity and evolution of rabies viruses [32]. To more completely understand the role of recombination in the evolution of Indian isolates of RABV, further studies using complete genome data are required.

Molecular phylogenetic analysis of both, N and G genes were performed using alignment-based and alignment-free methods. The RTD-based alignment-free method is based on frequency of k-mers and the relative order in which the k-mers occur in the sequences. The method is proved to be accurate and computationally efficient for analysis of gene and/or genome data as validated using a variety of human viruses causing NTDs such as Mumps, Rhino, Dengue and West Nile viruses [33, 34]. Regardless of the method used, the phylogenetic trees constructed using both N and G genes showed RABV isolates from this study belonged to two distinct linages namely Arctic lineage and Indian subcontinent lineage. This finding is in agreement with an earlier study that showed coexistence of two distinct lineages of RABV in India [13]. An earlier study reported that RABV isolates tend to form genetic clusters based on the geographical region within India [35]. However, the present study found that RABV isolates clustered based on membership to respective lineages rather than geographic proximity. Notably, this study reports that the RABV isolates belonging to Indian subcontinent lineage clustered closely with the isolates from Sri Lanka that belong to other subcontinent lineage. This highlights the continued incursion and/or circulation of the variant subcontinent lineages of RABVs in India that might have been originated from Sri Lanka. Phylogenetic analysis also revealed that the Arctic like RABV isolates from this study clustered closely with human RABV isolates from India and Germany.

Complex mechanisms shape the ability of rabies viruses to be maintained within its primary host species than to be serially transmitted to a new host species [36]. RABV causes lethal infection in dogs with an infectious period spanning less than 1 week (and typically only 2–4 days) and relies on transmission among members of the same species to be maintained at the population level [37]. Continued circulation of the two distinct linages of RABVs in India despite several vaccination and stray dog population control campaigns suggest that there is a perpetuated transmission among dogs and enzootic viral establishment of the two distinct lineages among the dog population.

Several studies investigating RABV biology highlighted that RABVs are sensitive to control measures [38-40]. The maintenance of RABV is suggested to be driven by an interaction between density-dependent transmissions and rabies-induced mortality [41-43]. However, alternative mechanisms including demographic structure and spatial structure have recently been suggested to generate observed epidemic cycles for RABV maintenance in a country [44].

Approximately 36% of the world’s rabies deaths occur in India each year, most of which take place when children come into contact with infected dogs. Government of India, is implementing a “National Rabies Control Program” approved during 12th five year plan (2012-2017) which aimed to prevent the human deaths due to rabies and to prevent transmission of rabies through canine (dog) rabies control. Dog population management (DPM) involves improving the health and well-being of stray or community dogs by vaccination, and reduce dog population size by routine birth control programs which can facilitate more effective rabies control [45]. Despite vaccination for many years, there was no downward trend in the RABV incidence in India and there is continued co-circulation of two distinct lineages as found by this study. Intentional or unintentional translocation of dogs between different geographical regions could potentially compromise natural or vaccine-generated barriers [46-49]. As the Indian subcontinent lineage RABV isolates from Chennai are closely related to viruses from Sri Lanka, this suggests the possibility of regular incursions of RABVs between the two countries. Hence, it is also important to check the cross-boundary moment of dogs and implement proper quarantine to prevent the spread of rabies.

## Acknowledgments

The authors would express their sincere gratitude to Dr. Elankumaran Subbiah (late) who played a key role in the conception and design of these experiments, who sadly could not be a co-author of this publication. The authors would also like to acknowledge the access to computational facilities at the Bioinformatics Centre, SPPU, supported by the Department of Biotechnology, Govt. of India.

